## Supplementary figures for "Nuclear and cytosolic J-domain proteins provide synergistic control of Hsf1 at distinct phases of the heat shock response"

### Figure 1 - Figure Supplement 1

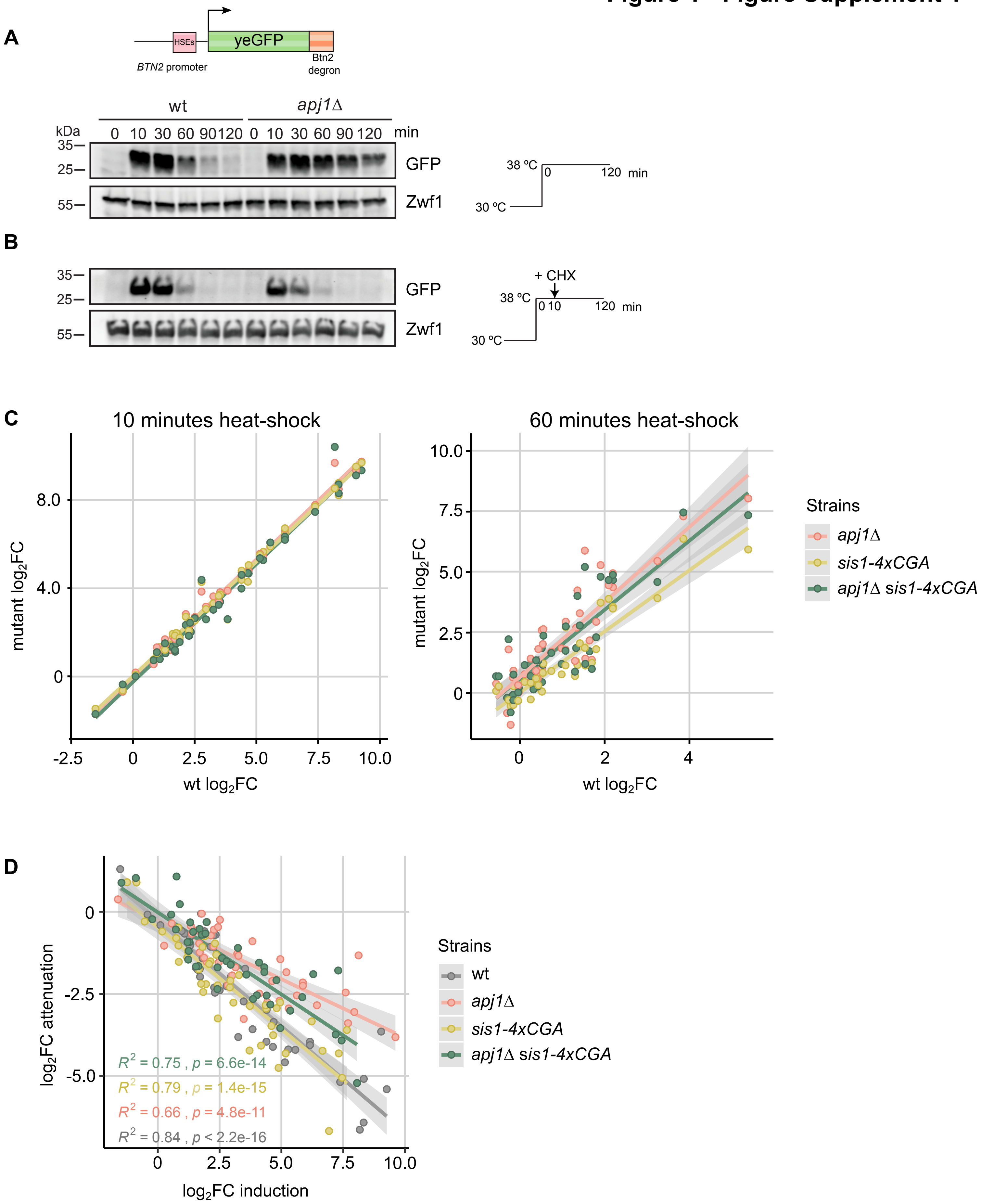

**Figure 3 - Figure Supplement 1**

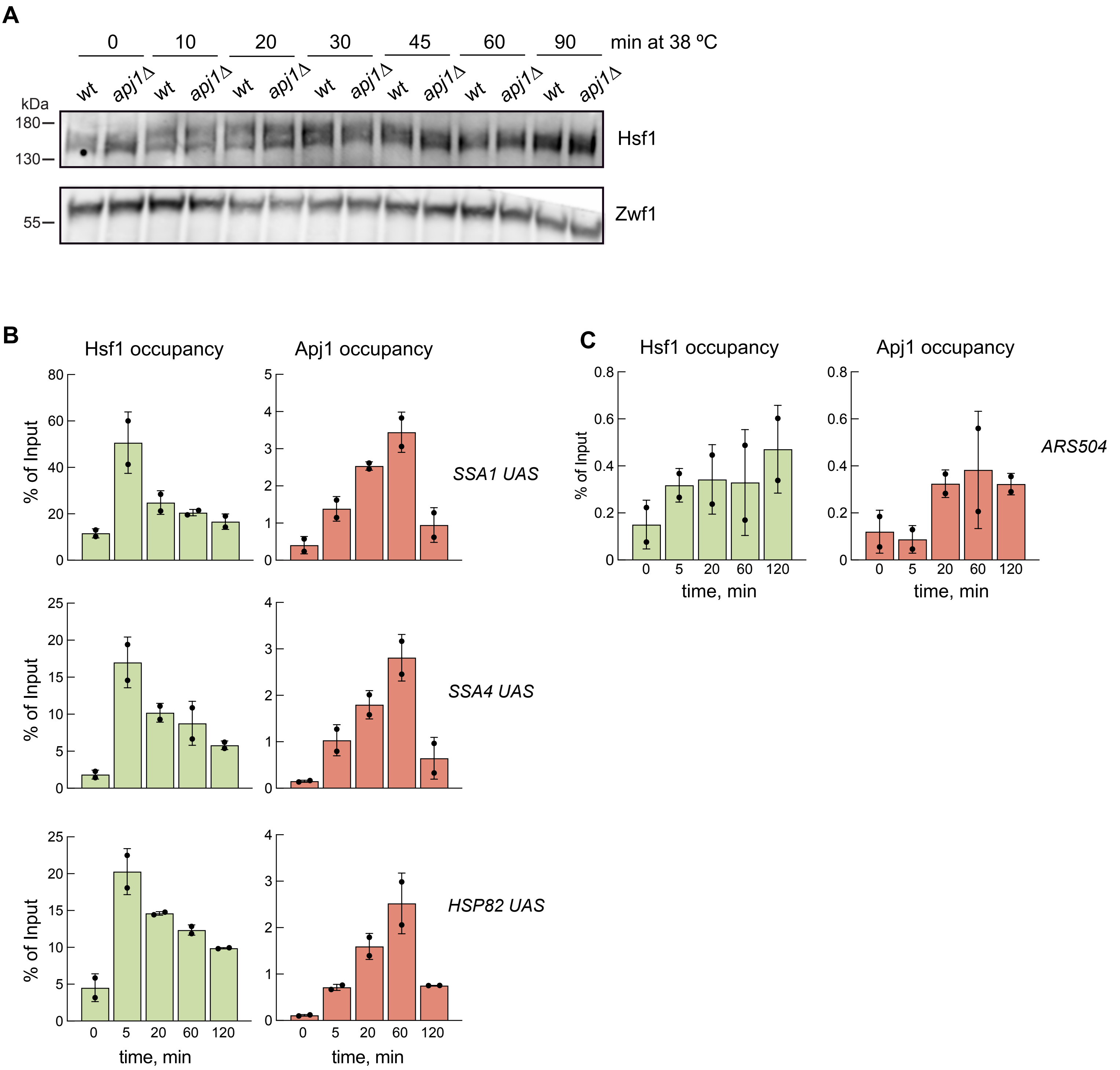

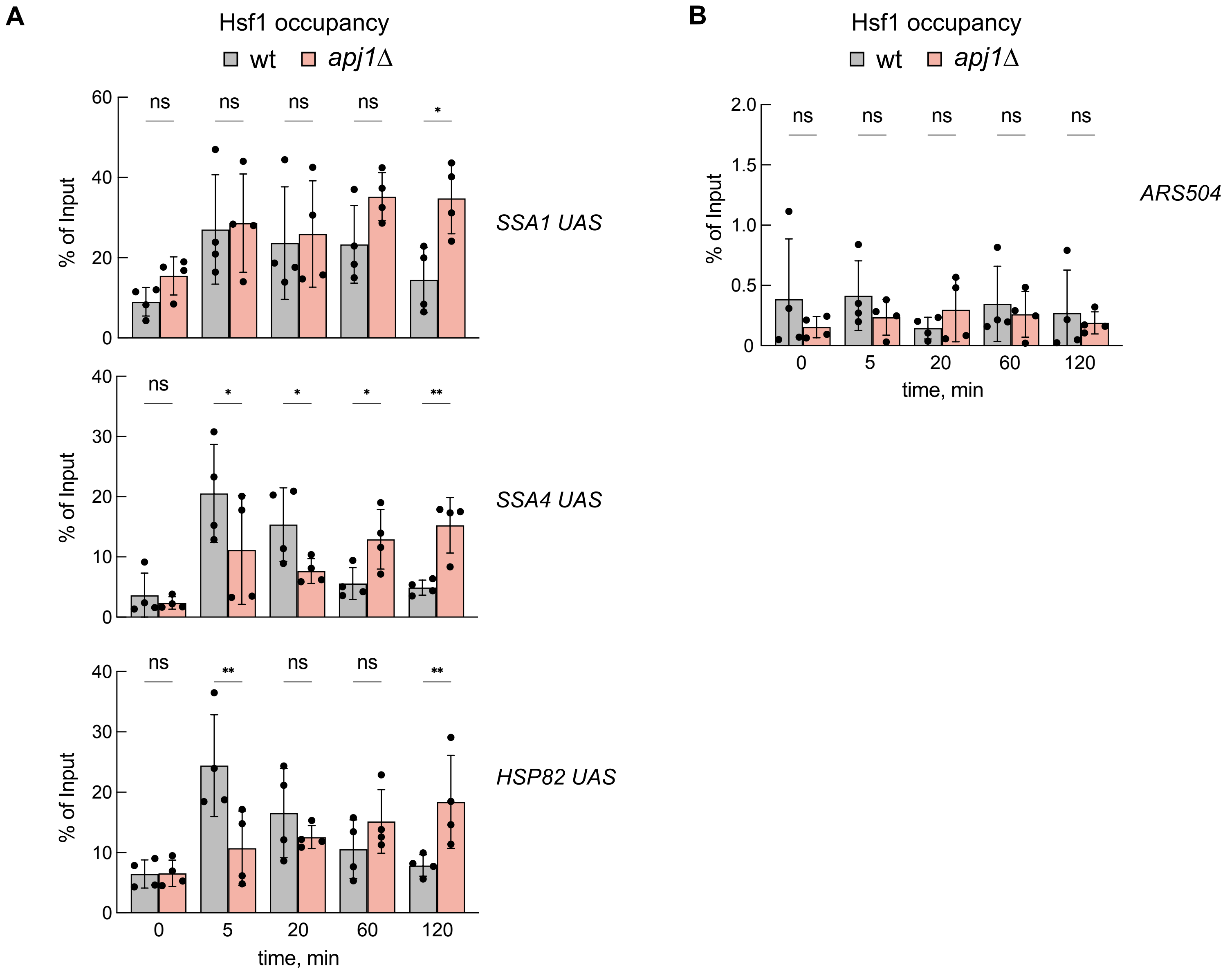


**Figure 4 - Figure Supplement 1**

### Figure 4 - Figure Supplement 2

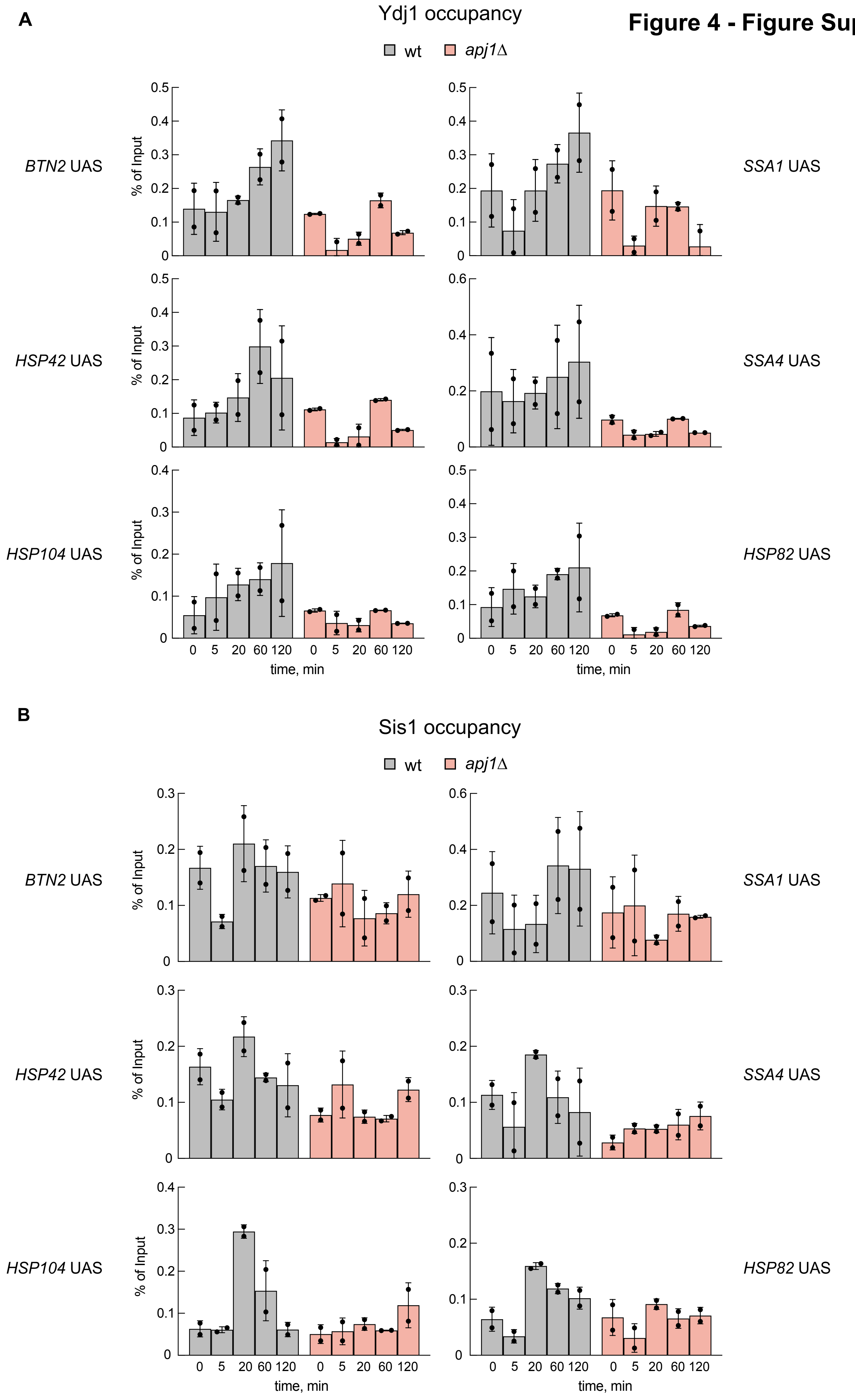

### Figure 5 - Figure Supplement 1

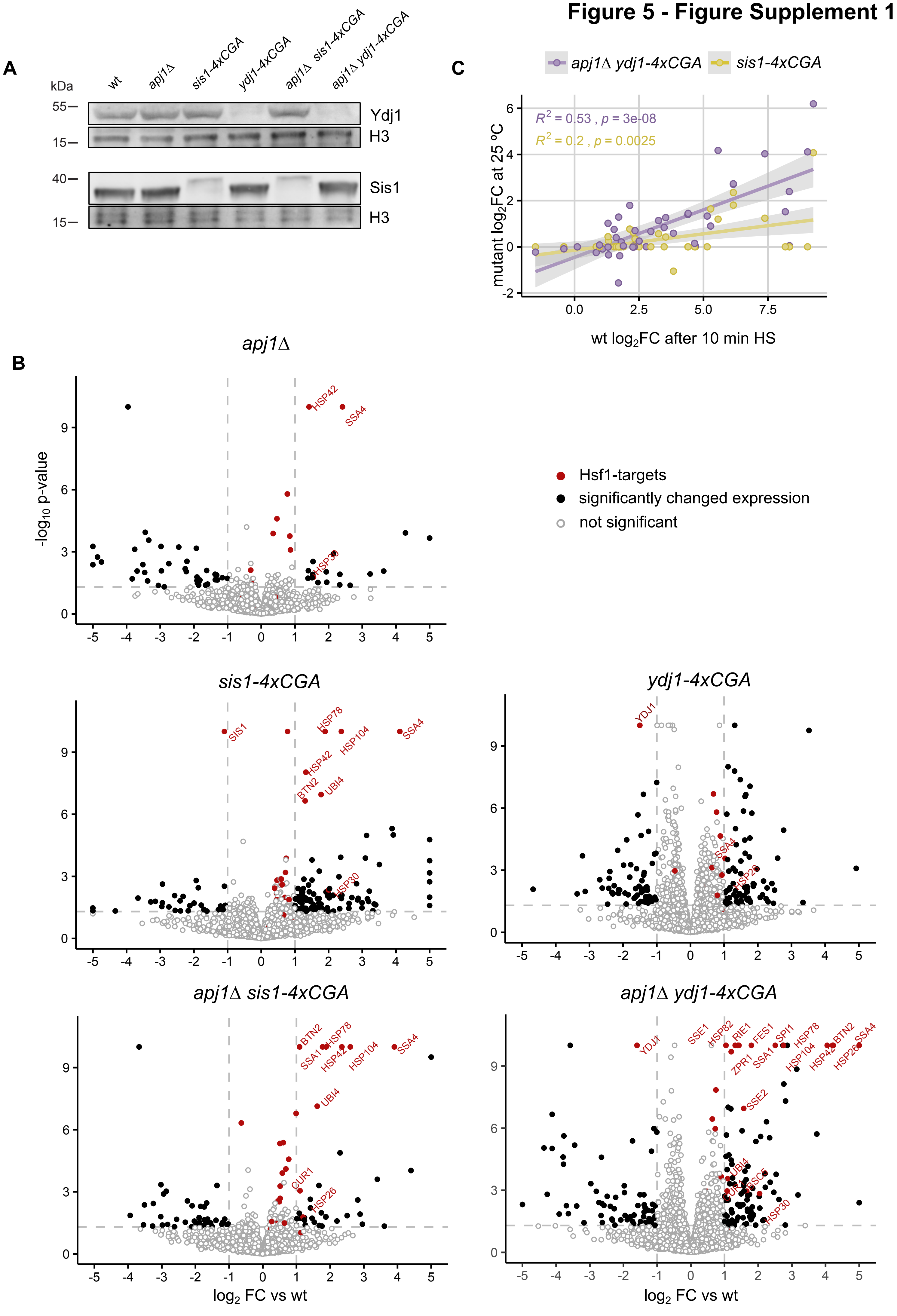

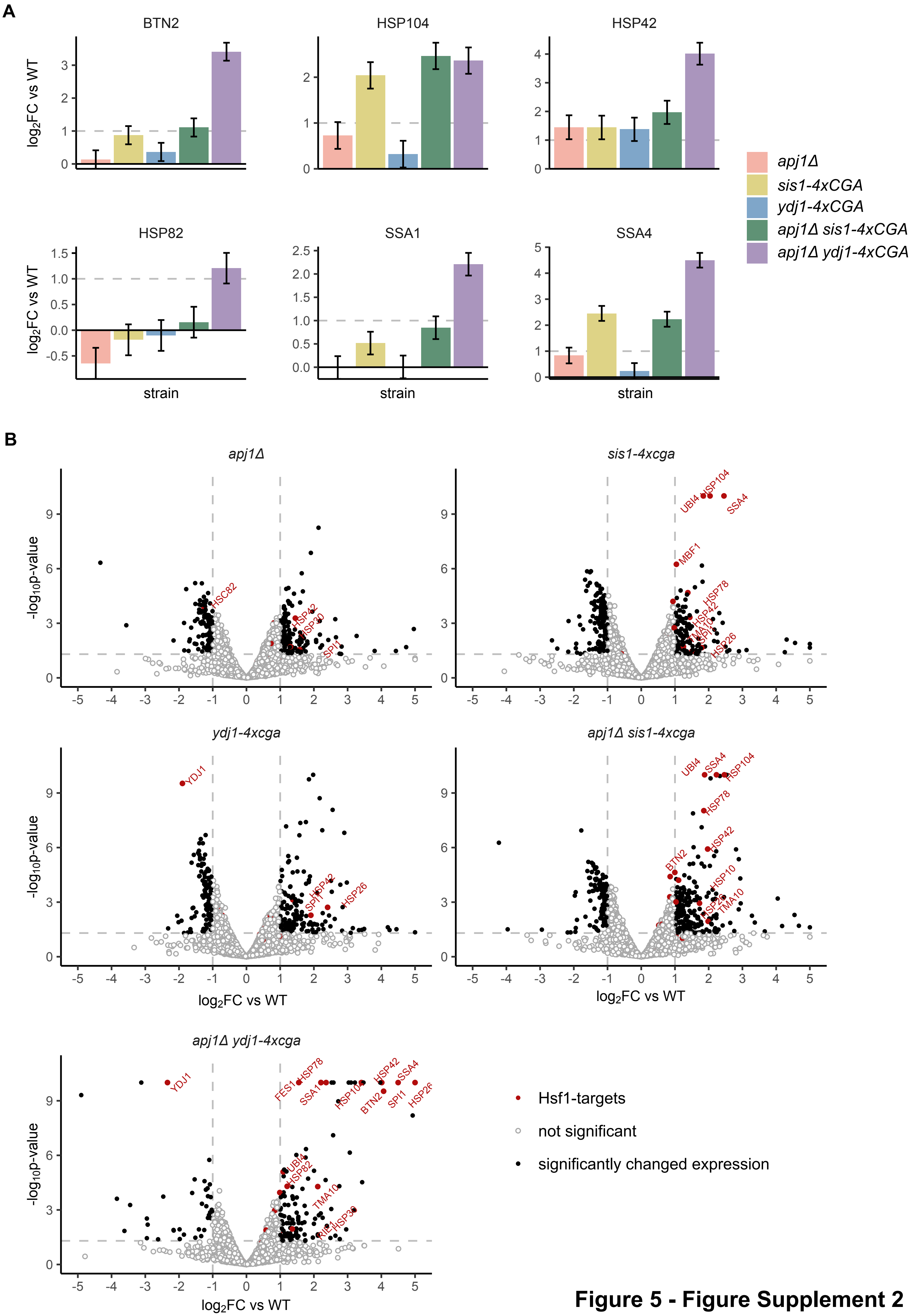


**Figure 5 - Figure Supplement 2**

**Figure 6 - Figure Supplement 1**

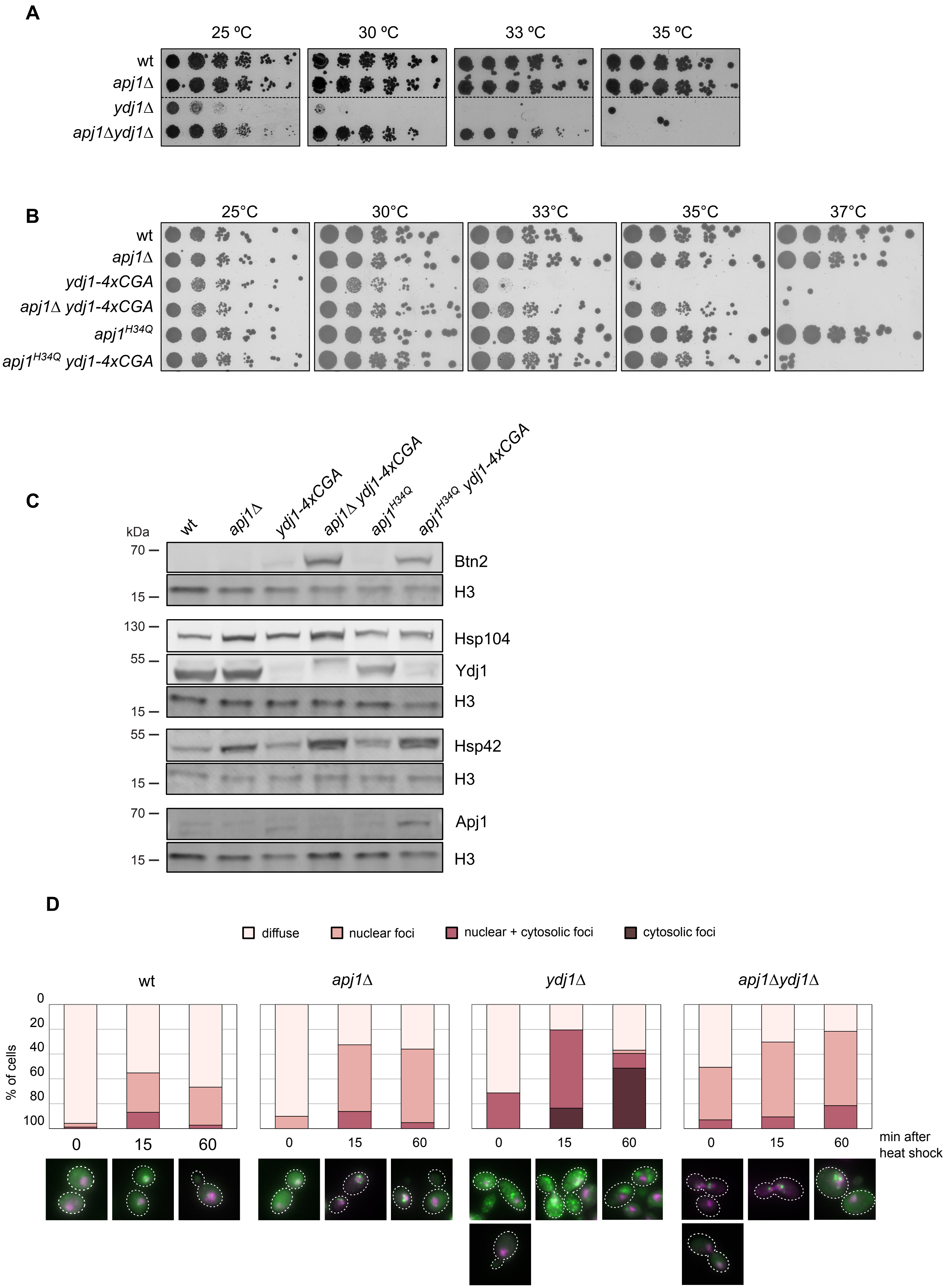

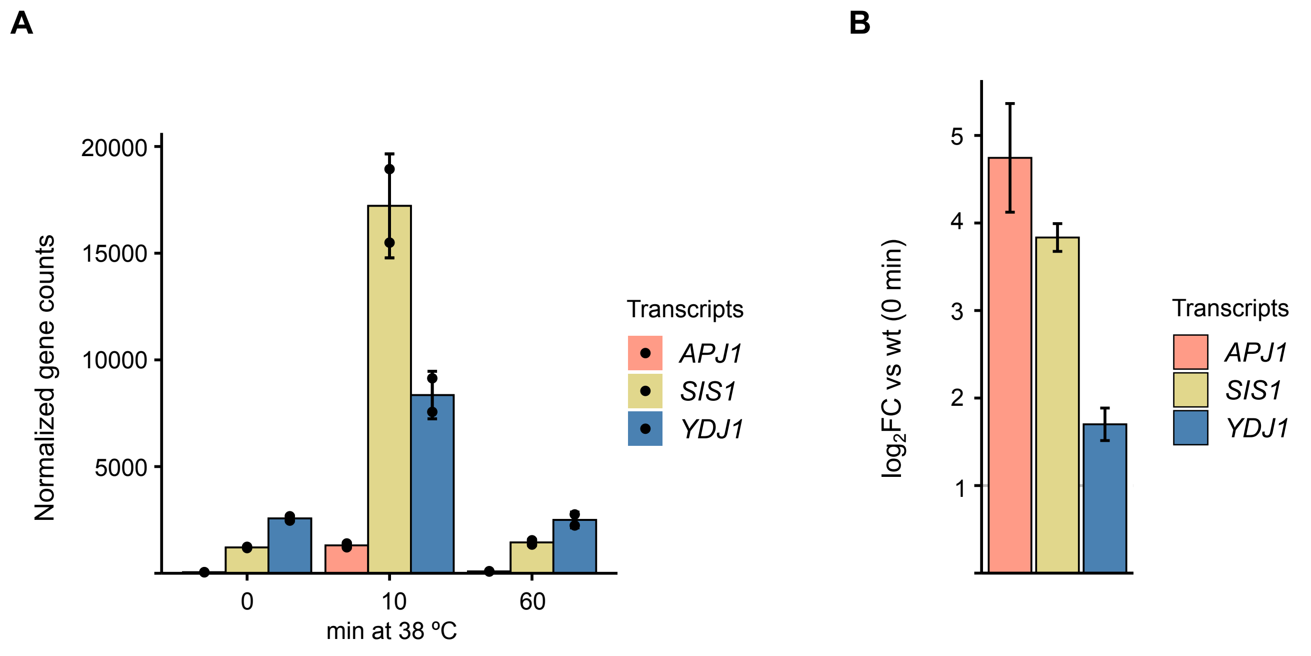


**Figure 1 - Figure Supplement 2**
