## Supplementary Tables S1-4 for "Nuclear and cytosolic J-domain proteins provide synergistic control of Hsf1 at distinct phases of the heat shock response"

| Name | Fenotype | Genotype | Source |
| --- | --- | --- | --- |
| BY4741 | wt | *MATa his3Δ1 leu2Δ0 met15Δ0 ura3Δ0* | EUROSCARF |
| W303 - 1B | wt | *MATα leu2-3,112 trp1-1 can1-100 ura3-1 ade2-1 his3-11,15* | EUROSCARF |
| ASMY09 | wt | BY4741 *pdr5Δ::natNT2* | this study |
| ASMY118 | wt | BY4741 *pdr5Δ::natNT2*, *BTN2*prom-yeGFP_Btn2NTD*::hphNT1::HIS3* | this study |
| ASMY121 | *apj1Δ* | BY4741 *pdr5Δ::natNT2*, *BTN2*prom-yeGFP_Btn2NTD*::hphNT1::HIS3*, *apj1Δ::KanMX4* | this study |
| ASMY144 | Hsf1-3xFLAG-V5 | BY4741 *pdr5Δ::natNT*2, *hsf1Δ::KanMX4, Hsf1pr-HSF1-3xFLAG-V5::HIS3 (C. glabarata)* | this study |
| ASMY156 | Hsf1-3xFLAG-V5, *apj1Δ* | BY4741 *pdr5Δ::natNT*2, *hsf1Δ::KanMX4, Hsf1pr-HSF1-3xFLAG-V5::HIS3 (C. glabarata)*, *apj1Δ::hphNT1* | this study |
| FA2216 | *hsf1-848* | BY4741 *hsf1-848::KanMX6, cir°* | this study |
| FA2220 | *hsf1-848, apj1Δ* | BY4741 *hsf1-848::KanMX6, apj1Δ::hphNT1, cir°* | this study |
| CRY063 | *apj1Δ* | BY4741, *apj1Δ::HIS3MX6* | this study |
| CRY066 | Hsf1-GFP | BY4741 Hsf1-yeGFP::*HIS3MX6* | this study |
| CRY068 | Hsf1-GFP *apj1Δ* | BY4741 Hsf1-yeGFP::*HIS3MX6, apj1Δ::hphNT1* | this study |
| CRY108 | Hsf1-GFP *ydj1Δ* | BY4741 Hsf1-yeGFP::*HIS3MX6, ydj1Δ::URA3* | this study |
| CRY110 | Hsf1-GFP *apj1Δydj1Δ* | BY4741 Hsf1-yeGFP::*HIS3MX6, apj1Δ::hphNT1, ydj1Δ::URA3* | this study |
| CRY104 | *ydj1Δ* | BY4741 *pdr5Δ::natNT2, BTN2*prom-yeGFP_Btn2NTD::*hphNT1::HIS3,ydj1Δ::URA3* | this study |
| CRY106 | *apj1Δydj1Δ* | BY4741 *pdr5Δ::natNT2, BTN2*prom-yeGFP_Btn2NTD::*hphNT1::HIS3, apj1Δ*::*KanMX4*, *ydj1Δ::URA3* | this study |
| *ASK804* | BUD3-HS | BY4741 *UASHS-BUD3* | Chowdhary, S. Kainth, A. et al 2019 |
| GMY008 | Apj1-13xMyc | BY4741 Apj1-13XMyc::*KanMX* | this study |
| GMY011 | Apj1-13xMyc, BUD3-HS | BY4741 UASHS-BUD3, Apj1-13xMyc::*KanMX* | this study |
| GMY019 | *apj1Δ* | BY4741 *apj1Δ::hphMX* | this study |
| YBK26 | *apj1Δ* | BY4741, *apj1Δ::hphMX, pdr5Δ::natMX* | this study |
| GMY062 | Sis1-13xMyc | BY4741 Sis1-13XMyc::*KanMX* | this study |
| GMY063 | Sis1-13xMyc, *apj1Δ* | BY4741 Sis1-13XMyc::*KanMX*, *apj1Δ::hphMX* | this study |
| GMY064 | Ydj1-13xMyc | BY4741 Ydj1-13XMyc::*KanMX* | this study |
| GMY065 | Ydj1-13xMyc, *apj1Δ* | BY4741 Ydj1-13XMyc::*KanMX*, *apj1Δ::hphMX* | this study |
| SMY343 | Sis1-GFP | BY4741 pdr5Δ::natMX4, Sis1-yeGFP::*HIS3MX6* | Bukau's lab collection |
| CRY032 | Sis1-GFP, *apj1Δ* | BY4741 pdr5Δ::natMX4, Sis1-yeGFP::*HIS3MX6, apj1Δ::hphNT1* | this study |

| | | | |
|---|---|---|---|
| CRY170 | Sis1-GFP, *ypj1∆* | BY4741 pdr5∆::natMX4, Sis1-yeGFP::*HIS3MX6, ydj1∆::URA3* | this study |
| CRY172 | Sis1-GFP, *apj1∆ydj1∆* | BY4741 pdr5∆::natMX4, Sis1-yeGFP::*HIS3MX6, apj1∆::hphNT1, ydj1∆::URA3* | this study |
| JTY001 | wt | W303, GFP-lacI::*HIS3, HSP12-lacO128::URA3, HSP104-lacO256::TRP1, SEC63-13xMyc::KanMX,* POM34-mCherry::*natMX* | Chowdhary, S. Kainth, A. et al 2019 |
| CRY088 | *apj1∆* | W303, GFP-lacI::*HIS3, HSP12-lacO128::URA3, HSP104-lacO256::TRP1, SEC63-13xMyc::KanMX,* POM34-mCherry::*natMX*, *apj1∆::hphNT1* | this study |
| CRY164 | *ydj1∆* | W303, GFP-lacI::*HIS3, HSP12-lacO128::URA3, HSP104-lacO256::TRP1, SEC63-13xMyc::KanMX,* POM34-mCherry::*natMX*, *ydj1∆::BleMX6* | this study |
| CRY166 | *apj1∆ydj1* | W303, GFP-lacI::*HIS3, HSP12-lacO128::URA3, HSP104-lacO256::TRP1, SEC63-13xMyc::KanMX,* POM34-mCherry::*natMX*, *apj1∆*::hphNT1, *ydj1∆::BleMX6* | this study |
| LSY009 | *ydj1-4xcga* | BY4741, ydj1-4xcga::NatMX | this study |
| LSY012 | *sis1-4xcga* | BY4741, sis1-4xcga::NatMX | this study |
| LSY020 | *apj1∆ ydj1-4xcga* | BY4741, apj1∆::HIS3MX6, ydj1-4xcga::NatNT2 | this study |
| LSY019 | *apj1∆ sis1-4xcga* | BY4741, apj1∆::HIS3MX6, sis1-4xcga::NatNT1 | this study |
| LSY072 | *hsf1-848 ydj1-4xcga* | BY4741 hsf1-848::KanMX6, cir°, ydj1-4xcga::NatNT2 | this study |
| LSY073 | *hsf1-848 apj1∆ ydj1-4xcga* | BY4741 hsf1-848::KanMX6, apj1∆::hphNT1, cir°, ydj1-4xcga::NatNT2 | this study |
| LSY49 | *apj1H34Q* | BY4741, *apj1H34Q::HIS3MX6* | this study |
| LSY067 | *apj1H34Q ydj1-4xcga* | BY4741, apj1H34Q::HIS3MX6, ydj1-4xcga::NatNT2 | this study |
| LSY074 | wt pRS315 EV | BY4741, pRS315 | this study |
| LSY075 | *apj1∆* pRS315 EV | BY4741, *apj1∆::HIS3MX6*, pRS315 | this study |
| LSY076 | *ydj1-4xcga* pRS315 EV | BY4741, ydj1-4xcga::NatMX, pRS315 | this study |
| LSY077 | *apj1∆ ydj1-4xcga pRS315 EV* | BY4741, apj1∆::HIS3MX6, ydj1-4xcga::NatNT2, pRS315 | this study |
| LSY078 | wt TDH3:Sis1 | BY4741 pRS315 TDH3:Sis1 | this study |
| LSY079 | *apj1∆* TDH3:Sis1 | BY4741, *apj1∆::HIS3MX6*, pRS315 TDH3:Sis1 | this study |
| LSY080 | *ydj1-4xcga* TDH3:Sis1 | BY4741, ydj1-4xcga::NatMX, pRS315 TDH3:Sis1 | this study |
| LSY081 | *apj1∆ ydj1-4xcga TDH3:Sis1* | BY4741, apj1∆::HIS3MX6, ydj1-4xcga::NatNT2, pRS315 TDH3:Sis1 | this study |

### Table S2:Plasmids

| **Backbone** | **Insert** | Source |
| --- | --- | --- |
| pRS315 | empty vector | Bukau's Lab collection |
| pRS315 | TDH3:Sis1 | Brandman's lab collection |
| pFA876 | pRS315 pApj1 GFP-Apj1-AAA | this study |
| pFA711 | pCU426 pGAL GFP | this study |

Table S3: Antibodies

### Table S3: Antibodies

| Antibody | Dilution | Source | |
|---|---|---|---|
| α-Btn2 (rabbit) | 1:5000 | Bukau's Lab collection | |
| α-Hsp42 (rabbit) | 1:5000 | Bukau's Lab collection | |
| α-Zwf1 (rabbit) | 1:50000 | Bukau's Lab collection | |
| α-Apj1 (rabbit) | 1:2000 | den Brave's Lab collection | |
| α-FLAG (mouse) | 1:10000 | Sigma-Aldrich | |
| α-H3 (rabbit) | 1:10000 | Sigma-Aldrich | |
| α-Sis1 (rabbit) | 1:5000 | Bukau's Lab collection | |
| α-Ydj1 (rabbit) | 1:5000 | Bukau's Lab collection | |
| α-Hsp104 (rabbit) | 1:20000 | Bukau's Lab collection | |
| α-GFP (rabbit) | 1:1000 | Bukau's Lab collection | |
| α-cMyc (mouse) | 1:360 | Santa Cruz Biotechnology | 2.5 µl of Ab per sample (ChIP) |
| α-Hsf1 | 1:600 | Gross' Lab collection | 1.5 µl of Ab per sampl (ChIP) |
| Alkaline Phosphatase Goat Anti-Rabbit IgG, AP1000 | 1:2500- 1:10000 | Vector Laboratories | |
| Alkaline Phosphatase Goat Anti-Mouse IgG, AP2000 | 1:2500- 1:10000 | Vector Laboratories | |

**Table S4: RT-qPCR Primer**

**Figure 3B**

| Primer Name | Sequence (5' to 3') |
|---|---|
| pTOS1_RT_for | ACCGACTAATGCGGTCATGGAAAGC |
| pTOS1_RT_rev | CTTTTCTCGCAAGAAGACTCCAGAATCA |
| pSSA4_RT_for | GGATATCTTTTGCCCGGTGAGTTG |
| pSSA4_RT_rev | TGTCGTCAAACTAAGGAGCTTCCC |
| pUBI4_RT_for | GGAGCATCACACAGCCGTACATC |
| pUBI4_RT_rev | AAAAGGAGGAACCGCCCTCAAATG |
| pSSA3_RT_for | GATGCCTATGGAGGTTATGGGTGC |
| pSSA3_RT_rev | CCCTTCCATTCGTTTCCAATTGTGC |
| pHSP42_RT_for | CACGCGCTTAAAAGTTCTGGAAGG |
| pHSP42_RT_rev | AACTAACTTCACAGAGGCCTCCCC |
| pBTN2_RT_for | GTGGAGCTCGAGAGTTGTATCCAG |
| pBTN2_RT_rev | CGCCAAGAACTGAAGGCTTCTATG |

**Figure 3D/E, Figure 4, Figure 3 - Figure Supplement 1, Figure 4 - Figure Supplements 1/2**

| | |
|---|---|
| ARS504 FP | GTC AGA CCT GTT CCT TTA AGA GG |
| ARS504 RP | CAT ACC CTC GGG TCA AAC AC |
| HSP104 UAS-267 FP | CTT AAA CGT TCC ATA AGG GGC |
| HSP104 UAS-216 RP | TGC AGT TCT TTG AGA TGG GCC |
| HSP82 UAS -394 FP | CCT CTC TCA ACA CAG TAA TCC ATA AAC |
| HSP82 UAS -242 RP | CTT CCA CGG CGT TCT AGA AAA AAA AG |
| SSA4 UAS -374 FP | GCC GCA CAT CCA TTC CGG TAT G |
| SSA4 UAS -312RP | CGG GCA AAA GAT ATC CGC TTT G |
| SSA1 UAS -428 FP | CGGTGTGTGGATGATGGTTTCATCAT |
| SSA1 UAS -178 RP | GTCCTCGAAACGATCAGCTAATCTAAATGG |
| HSP42 UAS -371 FP | GGATATGACATACTTCAATTCAGC |
| HSP42 UAS -160 RP | CAAGTCTTATATAACTAACTTCACAGAGG |
| BTN2 UAS -406 FP | GTCATGTAGCACTATTTCAGCC |
| BTN2 UAS -220 RP | CATTTGTTTTGCCACTTTACTTCG |
